# Circuit dynamics of superficial and deep CA1 pyramidal cells and inhibitory cells in freely-moving macaques

**DOI:** 10.1101/2023.12.06.570369

**Authors:** S. Abbaspoor, K.L. Hoffman

## Abstract

Diverse neuron classes in hippocampal CA1 have been identified through the heterogeneity of their cellular/molecular composition. How these classes relate to hippocampal function and the network dynamics that support cognition in primates remains unclear. Here we report inhibitory functional cell groups in CA1 of freely-moving macaques whose diverse response profiles to network states and each other suggest distinct and specific roles in the functional microcircuit of CA1. In addition, pyramidal cells that were segregated into superficial and deep layers differed in firing rate, burstiness, and sharp-wave ripple-associated firing. They also showed strata-specific spike-timing interactions with inhibitory cell groups, suggestive of segregated neural populations. Furthermore, ensemble recordings revealed that cell assemblies were preferentially organized according to these strata. These results suggest sublayer-specific circuit organization in hippocampal CA1 of the freely-moving macaques that may underlie its role in cognition.

## Introduction

Circuit dynamics specific to the hippocampus govern its role in navigation, memory, and social/motivational behaviors across species^1–7^. Based on studies in rats and mice, these dynamics arise from a well-described and diverse range of cell classes^8–11^. For example, the pyramidal cell layer is organized into radial-axis strata^12^, each receiving distinct afferents^13–16^, bearing different intrinsic and task-related response characteristics^15,17–23^, and projecting to different target structures^12,22^. The dynamics of these pyramidal cells are precisely regulated by inhibitory cell classes^10,24,25^ that differ from each other in their behaviorally specific firing patterns and roles in shaping network oscillations^11,26–30^. Because these inhibitory cell classes differently segregate pyramidal cell assembly patterns in space and time^29,31,32^ they may be fundamental determinants of the role of these assemblies in memory and navigation. More generally, the decisive and diverse roles of these cell types in circuit function make them critical targets in disease states such as in epilepsy, schizophrenia, and neurodegenerative diseases ^25,33–35^.

Although many aspects of hippocampal physiology are conserved between rodents and primates, differences in oscillatory dynamics^36–41^ and behavior-specific modulation^42–51^ suggest there may be phylogenetic specializations that may be uncovered by an understanding of the underlying functional cell composition. A first-pass bisection of hippocampal (and medial temporal lobe) cells in primates into putative inhibitory and excitatory classes (or four groups^52^) reveals characteristic responses at both the circuit-level^38,52–54^ and cognitive/behavioral levels^55,56^. Beyond that coarse division, we lack a description of primate CA1 functional cell groups, that can bridge what is known from anatomy and rodent physiology to circuit function and behavior. To understand the functional microcircuit in hippocampal CA1 of macaques, and its possible conservation or deviation from the canonical rodent CA1 region, we sought to identify functional cell groups, and their role in the local microcircuit function across task and sleep states. Additionally, we describe CA1 pyramidal cell activity in primates along their previously unexplored superficial and deep strata, including basic physiological responses, pairwise activity among the pyramidal groups and relative to putative inhibitory cell groups, and finally, in relation to network properties including oscillations and synchronized assemblies of activity.

## Results

### Identification of functional cell groups in laminar recordings of macaque CA1

We recorded from hippocampal CA1 layers in two freely-moving macaques, as they performed a sequential memory task and during sleep (Figure 1, A and B). Across 35 daily sessions (M1: 17 sessions, M2: 18 sessions) we measured the local field potentials and activity in ensembles of units. To align recording depths across these sessions and animals, we calculated for each session the current source density (CSD) during sharp-wave ripple (SWR) events. SWRs are known to evoke sinks in the stratum radiatum and sources in the stratum pyramidale layers of CA1^57^ (Figure 1C). Within this region, we found the slope of the slow component of sharp-wave ripples, which crossed zero across all sessions and animals, and we aligned to that zero-crossing. To validate that alignment, we measured ripple power, which peaks within the central region of the pyramidal layer (Figure 1C). After identifying single unit activity typically consisting of waveforms across several adjacent channels (Figure 1B), we selected the electrode contact with the largest spike amplitude for each unit as the best estimate of the location of the cell body. The regular linear distribution of these recording sites on the probe shanks allowed us to determine the relative depths of the cell bodies of the simultaneously recorded neurons (Figure 1D).

**Figure 1.**
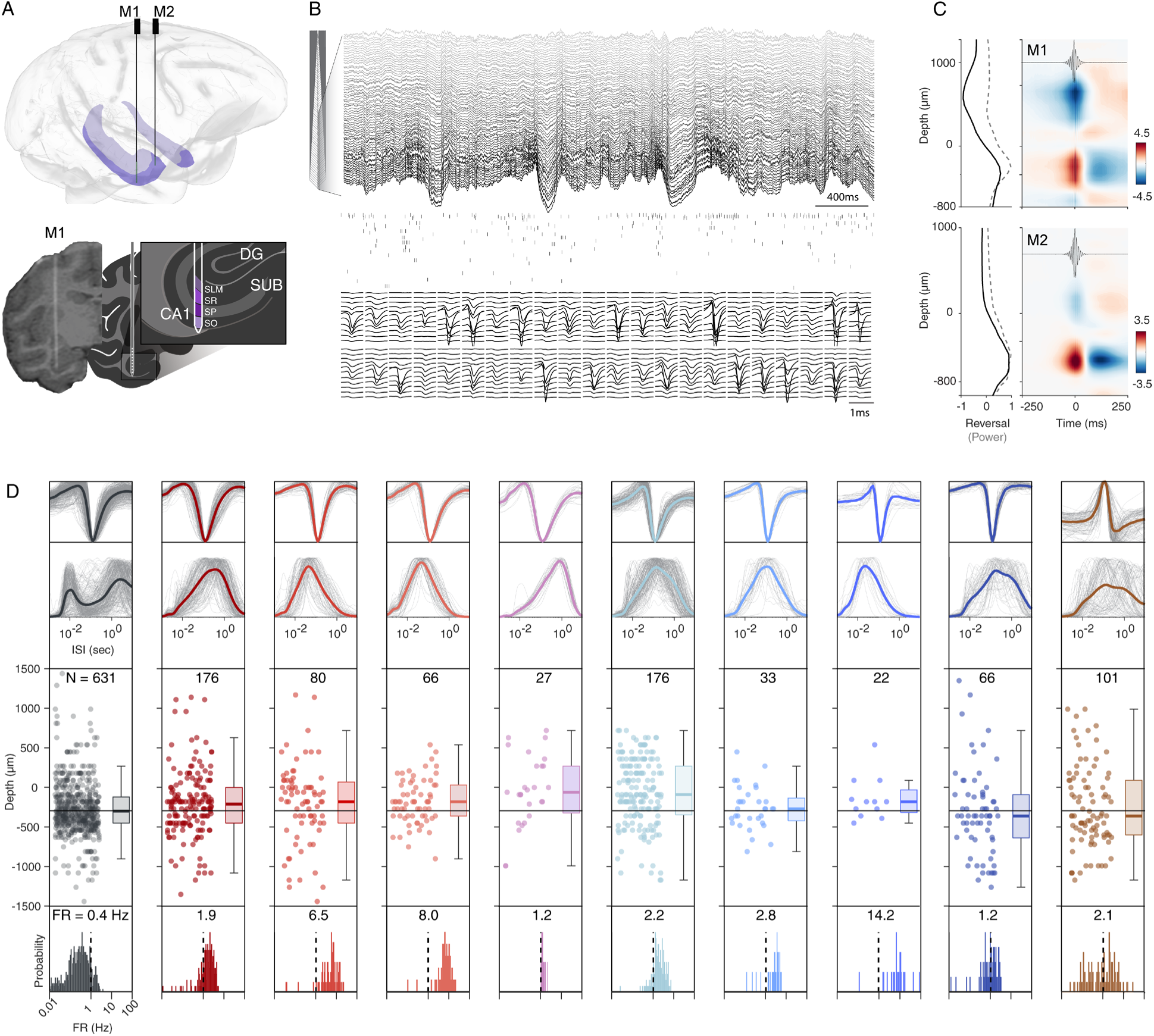
Single units from macaque CA1 characterized by depth and physiological parameters. **(A)** Top: schematic rendering of the electrode arrays localized to the target regions for each animal (M1, M2), in anterior hippocampal CA1. Dark purple: CA1. Both animals’ trajectories were localized with CT-MR registration and verified physiologically. Bottom: Coronal view of the CT coregistered to MR for one electrode trajectory, targeting the hippocampal CA1, and schematic of hippocampal CA1 layers. **(B)** Top: LFP traces across 64 of 128 recorded channels of one probe (40 micron spacing) spanning layers of hippocampal CA1. Middle: Spike raster of simultaneously recorded neurons from this probe (N = 48). Bottom: Example spike waveforms for selected units from this recording. **(C)** Top, left: Ripple slope reversal (black) and ripple power (dashed gray) across channels (depth). Top, right: Current source density of average ripple LFP for monkey 1 (blue: sink, red: source). Bottom: The same as top but for the 2^nd^ animal. **(D)** CA1 cell groups. 1^st^ Row: Grand mean (colored) and unit-mean (gray) waveforms (1.5 ms) for each cell group. 2^nd^ Row: Grand mean (colored) and unit-mean (gray) log interspike-interval distributions (0-10 s) for each cell group. 3^rd^ Row: Estimated depth of units aligned to ripple reversal (0 µm). Box plots (throughout) show median and quartiles. Depths differed by group, p < 0.001, Kruskal-Wallis test. Superficial-deep pyramidal split shown by black horizontal line (−330 µm). N= number of units per group. 4^th^ Row: Distribution of mean firing rates, shown for each cell group, dashed line at 1 Hz. Grand mean reported at top of plot. Firing rates differed by group, (p < 0.001, Kruskal-Wallis test).

We classified various cell groups with a semi-supervised approach according to their normalized waveform ^58,59^ and their interspike interval (ISI) distributions, to capture several of the principal intrinsic physiological characteristics that differ by cell group. This led to 10 separate cell groups (Figure 1D, see Methods; overall between-group distances exceeded within-group distances: p < 0.001, Kruskal-Wallis, Figure S1A). We use the term “cell groups” throughout to refer to these 10 clusters and note that these groups are clustered based on extracellular physiological features only, and thus may differ from the formal “cell classes” or “cell types” as identified through molecular/immunohistochemical methods or cytoarchitectonics.

The incorporation of the ISI distributions as features along with spike waveforms qualitatively drew out cells with bursting activity, low-firing rates, and broad waveforms, characteristic of CA1 pyramidal cells^60,61^. These comprised the first cell group (Figure 1D, far left, black, notably well-localized to the ripple layer i.e. Stratum Pyramidale). The remaining negative-deflecting groups were therefore considered putative inhibitory cells, because pyramidal cells are the only excitatory cell class in CA1^62^.

As expected from these features, firing rates of these cell groups differed (p<0.001, Kruskal-Wallis, Fig. 1D, Figure 1D and S1C inset). In addition, 3 neuron types selectively decreased firing during sleep compared to task (p<0.001, permutation test, Figure S1B). The ISI-conditioned coefficient of variation, CV2, measures the intrinsic variability of local spiking intervals, irrespective of global firing rate changes like those seen across behavioral epochs^63^. As expected, cell groups also showed differences in CV2 (p<0.01, Kruskal-Wallis test with post-hoc permutation test FDR corrected, Figure S1C), leading to distinct, cell group-characteristic joint ISI histograms (Figure S1D).

### Spectrolaminar profiles and spike-phase coupling by cell group

To better understand oscillatory composition across layers, we computed aperiodic-corrected power spectra (Figure 2A). Despite inter-subject differences in absolute peak frequencies across the spectrum, both subjects showed reduced theta-band power (5-10 Hz) during the task in comparison to early sleep overnight, across all recorded channels (p<0.05, cluster-based permutation test, Figure 2A), in line with previous studies^36–38^. Furthermore, the pyramidal layer showed relative increases in power in the mid frequencies of 15-40 Hz (Figure 2A), and each animal had a preferred higher gamma band apparent during the task (Figure S2A). These spectral profiles were also evident in the spike-phase coupling across units, measured as the pairwise phase consistency (PPC; Figure 2B and S2B). Comparing across states, spike-field coherence within the theta frequency range (5-10 Hz) was greater during sleep (p<0.05, two-sample permutation test) across all cell groups (Figure S2B). Conversely, spike-field coherence in the beta2/low (25-35 Hz) and higher (50-75 Hz) gamma ranges were significantly higher during the task (p<0.05, two-sample permutation test, Figure 2B), across cell groups.

**Figure 2.**
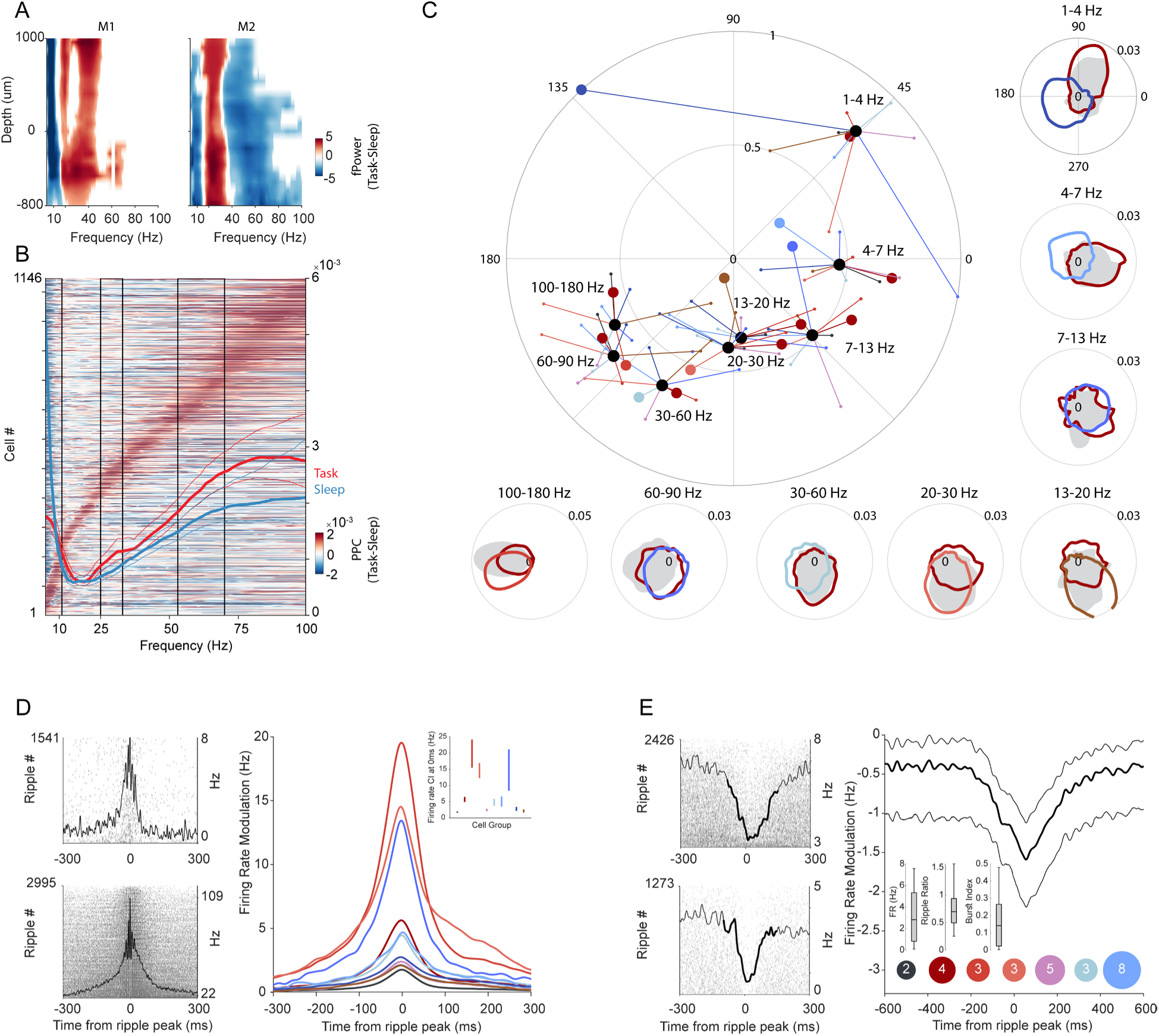
Local field potential dynamics and spike-field relationships across cell groups. **(A)** Spectrolaminar differences by behavioral state for each animal. Left: Color difference plot of 1/f corrected power spectrum of alert, task (red) and sleep states (blue) across depths of recordings for monkey 1. Non-significant differences by state are masked in white (significance set to p < 0.05, cluster-based permutation test) Right: Results for monkey 2 using the same conventions as for Monkey 1. **(B)** Pairwise phase consistency (PPC) differences between task and sleep states per unit, sorted based on each unit’s maximum PPC frequency. Mean PPC (thick lines) and bootstrap 95% CI (thin lines) are shown for the task in red, and for sleep in blue, with values shown on the right axis. Black boxes indicate those frequency ranges that differed across states (p<0.05, two-sample permutation test with Tmax multiple comparison correction). **(C)** Center, large: Phase-of-firing plot depicting the average resultant vector lengths and phase angles for each cell group, as a function of LFP frequency bands. Each color dot indicates the grand mean resultant vector length and angle for the respective cell group, plotted separately for each frequency band. Thick black dots represent the mean of those cell groups means, for each frequency band. Thick color dots represent the selected examples plotted on the off-center plots. Off-center, small: Normalized polar histograms showcasing the phase-of-firing probabilities for selected single neurons, organized by LFP frequency bands. Examples are colored according to group membership; gray shaded area shows examples from group 1 pyramidal cells. **(D)** Peri-SWR enhanced unit responses. Left: Example raster plot of individual neuron activity during ripple, and the superimposed line showing the PETH. Right: Lines show mean + bootstrap 95% CI for baseline-corrected ripple-associated activity for ripple-activated neurons (p < 0.05, one-sample randomization test) separated and color-coded according to cell groups. **(E)** Peri-SWR suppressed unit responses. Left: Example rasters as in (D), with the time window of significant modulation indicated by the bold black line. Right: conventions as in (D), for all ripple-suppressed cells. Inset: Quantile distributions across cells are shown for firing rate, ripple ratio and burst index (see Methods). Colored circles indicate the group identities and count of cells comprising the suppressed population.

Focusing only within band-limited bouts of high power, cells showed a generally large bandwidth, showing significant phase-locking to, on average, 4.4 of the 8 frequency bands. The grand mean phase angle per cell group revealed that these groups typically clustered within a limited range of preferred phases (∼90 degrees) that shifted by frequency (Figure 2C and S2C), but nevertheless differed reliably in mean phase-of-firing values for all but the lowest frequencies (except for 1-4 Hz, 4-7 Hz, and 7-13 Hz, p < 0.001, multi-sample Watson-Williams test, Figure S2C). Using the pyramidal cell group as a reference, the other cell groups deviated in phase angle as a function of frequency and cell group (Figure S2C-D; p < 0.05, two-sample Watson-Williams). Among cells exhibiting significant phase locking (p<0.001, Rayleigh test) within each frequency group, the largest resultant vectors were found in the following order: 1-4 Hz; 100-180 Hz and 4-7 Hz; 60-90 Hz; 30-60 Hz and 20-30 Hz; and finally, 7-13 Hz and 13-20 Hz; (p<0.05, Kruskal–Wallis test with Tukey-Kramer correction). Thus, the power spectral peaks were not necessarily the frequencies yielding the strongest phase-locking, nor the most consistent differences in phases of fining by cell types.

### Cell group-specific ripple-associated activity

Most cells increased their firing during ripples, across all cell groups (p<0.05, one-sample randomization test, Figure 2D and 3A) though to differing degrees (ripple ratio, during/baseline FR,p < 0.001 Kruskal-Wallis test, Figure S3B), and with differences in participation rate across ripple events (p<0.001 Kruskal-Wallis test, Figure S3B). The pyramidal cell group showed the highest ripple ratios but the lowest participation rate (p<0.001, pairwise permutation test, FDR corrected, Figure S3B). Putative inhibitory cell groups showed a striking range of participation probability and timing: some groups’ members participated in all ripple events (Figure S3B), and some restricted firing to a specific time relative to the ripple peak (Figure S3B; p < 0.001, Bartlett’s test). A small population of cells had suppressed activity during the ripple, and these were comprised mainly of the inhibitory (non-pyramidal) cell groups (p < 0.05, one-sample randomization test, Figure 2E).

**Figure 3.**
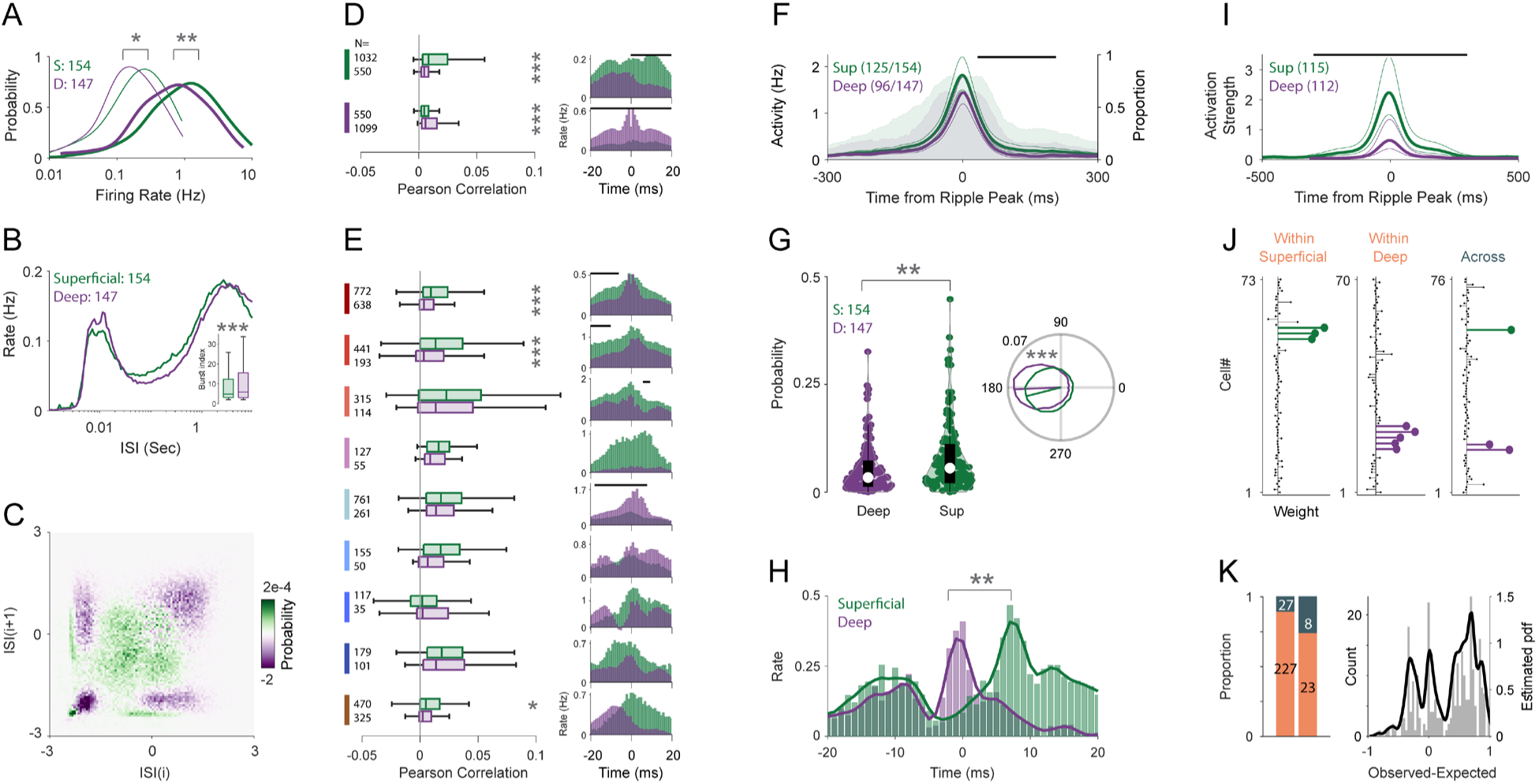
Sublayer-specific circuit dynamics of hippocampal pyramidal cells in macaque CA1. **(A)** Overall (thin lines) and ripple (thick lines) firing rate distribution for superficial (green, N= 154) and deep (purple, N= 147) pyramidal cells. **(B)** Mean ISI distribution. Inset: burst index distribution. **(C)** Difference (Sup-Deep) plot of joint probability density of current ISIs versus next ISIs. **(D)** Left: Pairwise cofiring of pyramidal cells with other pyramidal cells from within and across Sup, Deep groups. Referent group of pair shown by color code bar on the far left; target groups are Sup above (green), and Deep (purple), immediately below. Right: Mean cross-correlograms of sup/deep pyramidal cells. Black horizontal lines above plots (throughout) indicate significant difference (p<0.05, two-sample permutation test with Tmax multiple comparison correction). CCG values were baseline corrected (mean from −50 to −20 ms was subtracted from all bins). **(E)** Left: Pairwise cofiring of sup/deep pyramidal cells and putative inhibitory cell groups. Conventions as in (D). Mean cross-correlograms of sup/deep pyramidal cells and inhibitory cells. Conventions as in (D). **(F)** Ripple-aligned spike density (mean + 95% CI) for significantly modulated cells (N = 125 superficial, and 96 deep), black horizontal line shows significant differences (p < 0.05 cluster-based two-sided permutation tests). The proportion of significantly modulated units relative to the ripple peak is shown in the shaded area plots, units on the right axis (p<0.05, one-sample cluster-based permutation test). **(G)** Left: Distribution of the ripple participation probability. Right: Phase concentrations and mean resultant vector for significantly phase locked units (N = 74 superficial, 98 deep, p = 1e-3 Kuiper two-sample test/p=2e-4 permutation test). **(H)** Mean cross-correlograms of sup/deep pyramidal cells and ripple-suppressed inhibitory cells (N= 16 superficial, 20 deep,p < 0.05, two-sample permutation test with Tmax multiple comparison correction). **(I)** SWR-triggered average activation strength in post-task sleep for Deep (N = 115) and Sup (N = 112) cell assemblies. (p<0.05, two-sample cluster-based permutation test) **(J)** Example assembly weights for two “within” assemblies and one “across”. Rows indicate cells, and their length indicates their weight in that assembly. Significant members are colored. **(K)** Left: Fraction of within (N =227, orange) and across (N=27, dark gray) assemblies for all sessions and assemblies (N = 254) and for balanced sessions where the number of recorded deep and superficial were equal and assemblies that had at least 2 significant pyramidal members (N = 31). Right: Difference between expected proportions of within/across and observed proportions (N = 254). For a, b, c, d, e, and g, P values were computed using a two-sided permutation test (N = 5000).

### Stratification into superficial and deep CA1 pyramidal cells reveals physiological heterogeneity

To extend findings from the rodent literature, we examined whether CA1 pyramidal cells recorded in different depths of Stratum Pyramidale differ in their spiking properties and circuit-level dynamics. For each session, cells in the ‘Pyramidal’ group (meeting firing rate, burst and depth criteria), were median split according to depth, creating the CA1sup group (closer to Stratum Radiatum) and the CA1deep group (closer to Stratum Oriens). Compared to CA1deep cells, the firing rates of CA1sup cells were higher overall and during ripples, and they had a greater CV2 (P < 0.05, permutation test; Figure 3A-3C, Figure S1C), though they were less ‘bursty’ than their deeper peers (P < 0.001, permutation test; Figure 3B).

To investigate whether these cells interact differently, either within their subgroups, or in their coupling with the inhibitory groups, we analyzed their pairwise co-firing statistics. We found that pyramidal cells within a sublayer were more likely to fire together compared with cells across sublayers (Figures 3D), and they showed differences in co-firing strength with several of the inhibitory cell groups (p<0.001, permutation test; Figure 3E). The finer-grained temporal dynamics were evaluated using baseline-normalized cross-correlograms (CCGs). This revealed preferential firing within CA1sup and CA1deep (p<0.01, two sample permutation test with Tmax correction, Figure 3D). Furthermore, time lags between pyramidal and inhibitory cell pair spikes depended on both pyramidal cell strata and inhibitory cell group (p<0.01, two sample permutation test with Tmax correction, Figure 3E). Finally, CA1sup and CA1deep cells have somewhat opposite timing interactions with the ripple-suppressed cells: for CA1deep cells, the interactions peak at 0 ms, when CA1sup cell interactions are near their nadir; CA1sup cells’ peak is at approximately 7ms lag (p<0.01, two sample permutation test, Figure 3H).

During ripples, CA1sup had a higher participation probability and fired more than CA1deep (p<0.01 two-sample permutation test, Figure 3G). Relative to pre-ripple firing, CA1sup cells fired more post-ripple compared to CA1deep (p<0.05 cluster-based permutation test, Figure 3F). Finally, CA1deep cells are more strongly locked to the ripples and fire at a slight but significant phase advance compared to CA1sup (p<0.01 permutation test, p < 0.001 Kuiper’s test, Figure 3G).

### CA1 cell assembly membership by strata and cell group

We used unsupervised detection of recurrent co-activity to identify cell assemblies during sleep epochs, using established methodologies (Figure 3J and S4A, Methods). Assemblies that included pyramidal cell contributions were assigned to “within superficial,” “within deep,” or “across”, based on the category membership of those pyramidal cells. Our findings revealed that “within CA1sup” assemblies exhibited significantly stronger activation during ripple events in comparison to “within CA1deep” assemblies (p < 0.05 cluster-based permutation test, Figure 3I). Furthermore, assembly membership depended on layer, with most having all members belonging to the same sublayer (Figure 3K). Importantly, these results held after accounting for the priors (the observed exceeded expected sampling distributions of eligible cells by layer), and when only considering assemblies detected during balanced sessions, where an equal number of CA1sup and CA1deep cells were recorded, and assemblies had more than 2 significant pyramidal cell members (Figure 3K).

A cell’s selectivity across assemblies can be defined as the inverse of its participation rate (the proportion of assemblies to which a neuron significantly contributed, divided by all detected assemblies in a session). The majority of cells demonstrated a propensity for participating in specific assemblies, with a mean participation rate of 7% (Figure S4B). Participation varied by cell group (Figure S4C, p < 0.001, Kruskal-Wallis test and p < 0.05; post-hoc pairwise permutation test with FDR correction), and there was an overall negative correlation between the firing rate of neurons and their participation rate (p < 0.001, t-test Figure S4D). That is, inhibitory cell groups, overall, were more selective in their cell assembly participation than pyramidal cells.

## Discussion

In this study, we sought to forge a link between the extensive understanding of CA1 microcircuit function from rats and mice ^64,65^ to the network dynamics and brain-behavior states seen during naturalistic experiences in primates ^36,37,42,66,67^. Building on previous work ^58,59,68^, the semi-supervised clustering of waveforms and ISI distributions of isolated CA1 single units identified functional cell groups with diverse physiological characteristics. These included a range of firing patterns overall and in response to specific states of the local microcircuit including during sharp-wave ripples and across awake to sleep states. The phase relationship between the putative pyramidal cells and the inhibitory cells also varied as a function of network oscillations, suggesting dynamic interactions between these cell groups that may underlie the generation of these oscillations. Within the CA1 pyramidal cell group, we found physiological and functional segregation between those located closer to the stratum radiatum (‘superficial’) and those located closer to stratum oriens (‘deep’). These superficial and deep cells showed robust differences in their firing rate, burstiness, and sharp-wave ripple-associated firing. Importantly, the segregation of pyramidal cells into sublayers uncovered distinct cofiring patterns as a function of the depth of the pyramidal cells and the reference inhibitory cell group. Consistent with this notion, superficial and deep pyramidal cells preferentially formed sublayer-specific CA1 cell assemblies.

### Cell group classification

The taxonomy of neurons into cell types is foundational to understanding the organization of functional and dysfunctional circuit dynamics ^69,70^. Neuronal cell types can be identified through morphology, molecular composition, transcriptomics, connectivity, and physiological traits ^71,72^. The use of transcriptomic labeling and optical monitoring is typically limited in primates by factors like the inefficacy of viral serotypes ^73^, the need for optical access and head-fixation in vivo ^74^, the depth of relevant neural populations, and the number of simultaneously monitored cell types ^75,76^. As a consequence, we know less about cell types in deep structures, including the hippocampus, for those species that bear special relevance for understanding human disorders^77^. Furthermore, emerging evidence demonstrates that transcriptomic cell type taxonomies alone may not be sufficient to capture meaningful differences in cell types, particularly in light of species differences in physiology and morphology between primates and rodents ^78,79^. To improve the classification of neuronal cell types and the functional characterization of single neurons, multimodal datasets incorporating electrophysiological properties are essential to refine and extend foundational cellular taxonomies of the brain ^78,80^. In addition, an advantage of extracellular recordings, from single electrodes to high-density probes, is that they can capture a broad range of neuron populations including those inaccessible to genetic or optical methods, offering scalability to any cortical area ^81–83^ with high sampling resolution. In unimodal electrophysiological datasets, action-potential waveform and firing patterns offer effective markers of functional cell-type diversity ^68,84^. Although, and indeed because, functionally-defined cell types may not match precisely the same types identified through histological and molecular features, quantitative approaches toward cell-type classification using extracellular features facilitate the discovery of principled diversity in neuronal dynamics. This is made possible by reducing the complexity of the organization of cells into their effective, functional primitives, and enabling their reproducible identification. As such, we have focused here on the physiological classification in macaques due to its emergent feasibility and its direct and immediate significance on circuit function in behaving animals.

A prevalent approach to cell classification in primates relies on binary classification based on waveform features ^55,85–88^. Binary classification is limited in fully capturing the structural and transcriptomic diversity in monkeys and mice ^9,89^, and recent findings challenge the adequacy of this approach ^90–98^. In addition to refinements to waveform classification^59,90,99,100^, cells’ firing patterns may prove vital for more accurate and reliable classification ^101–103^. The firing patterns and timing of neuron activity summarized by the inter-spike interval (ISI) distributions have previously been applied to differentiate inhibitory from excitatory cells across brain regions and species ^52,60,66,104–109^. Informed by prior studies ^58,68^, our method additionally merges unsupervised analysis of spike waveforms and firing patterns (ISI distributions) with supervised refinements, to maximize the use of available information in extracellular electrophysiology studies for cell group classification. One caveat to this approach is that unsupervised methods, which excel at capturing diversity, do not automatically ensure consistency across studies. Thus, future research should investigate how physiological diversity reported using nonlinear unsupervised methods co-vary with morphological and transcriptomic features on a cell-by-cell basis to understand the relationships among these three modalities as they pertain to cell-type definition or cell state-dependent variations (see ^103,110^). This will enable a recalibration of classification criteria in accordance with the emerging differences in morphology and physiology across species and brain structure.

To separate putative pyramidal from inhibitory cells, we relied on criteria established in previous electrophysiology and anatomy studies of macaque or rodent CA1^65^, that pyramidal neurons have 1) a characteristic sparse burst spiking activity ^61,111–114^, 2) low firing rates ^60,61^, and 3) dense localization within in the pyramidal layer ^62,113^. Because pyramidal cells are the sole reported excitatory type in the hippocampal CA1 ^62^, we classified other cell groups with negative polarity as putative inhibitory. Although some inhibitory cell types may engage in bursting activity depending on their input, the firing rates of these classes are notably higher than those of pyramidal cells ^104^. Furthermore, the slow-spiking GABAergic interneurons in the hippocampus (e.g. Ivy interneurons ^91^) lack bursting activity ^91^. Therefore, our multimodal criteria based on firing rate and burst index aimed to exclude inhibitory groups from being misclassified as pyramidal (lowering false positives). The resulting 8 distinct inhibitory groups should not be viewed as discrete categories, i.e., with sharp statistical borders. Instead, they represent the most self-consistent types of responses embedded within a much broader distribution of responses, consistent with what has been proposed in transcriptomically-defined cell types ^8^. Furthermore, the methods used in this study are not well-suited for determining the relative proportions of cells between groups. For example, although GABAergic local circuit inhibitory interneurons constitute only 10–15% of the total neuronal population in the rodent hippocampal CA1^25,64^, the proportion of classified inhibitory cells in the current dataset was higher. This discrepancy may be due to several factors including uneven sampling across layers by the linear recording arrays, differences in proportions of cells excluded or missed by e.g. firing rate criteria, and potential differences across species.

We also observed a group of positive-polarity units (Punits), constituting 7% of the total population, characterized by a positive peak in the waveform recorded at the electrode with the largest waveform amplitude. Recent findings suggest that these positive spikes near the soma correspond to return currents^115^. In the CA1 region, these Punits show place fields, phase precession, and phase-locking to ripples ^115^. Our findings also show that despite a multimodal inter-spike interval (ISI) distribution indicating heterogeneity in this group, these units still display sharp-wave ripple-associated firing, phase modulation, and distinct co-firing patterns with the pyramidal cell group.

Finally, in the neocortex, inhibitory and pyramidal neurons exhibit significant interspecies differences in structure ^116^, gene expression ^117^, prevalence and connectomics ^79^ and intrinsic physiological properties such as input resistance, firing threshold, and afterhyperpolarization ^118–120^. Consistent with these findings, the firing rates in homologous brain areas vary markedly between species, although firing regularity is maintained ^121^. These observations suggest that the transcriptomic and physiological profiles of primate cell types may not closely match those found in rodents. This further highlights the importance of studying cell types in primates and implies that these differences could underlie variations in cognitive abilities ^122^.

### Cell-group specific spike-field relationships

Extensive research on the organization of rodent CA1 has identified state-dependent roles for specific functional cell types in regulating synaptic dynamics, network oscillations, and cell assembly dynamics ^24,25,123,124^. Despite some degree of mismatch in hippocampal brain-behavioral states across rodent and primate clades ^36,42,44,66,67^, it is nonetheless tempting to identify possible correspondences between the present groups and identified cell classes in rodent CA1. During sharp-wave ripple (SWR) events, most cell groups exhibited increased firing rates, however, inhibitory cell groups showed greater participation probabilities but lower ripple modulation. This indicates that pyramidal cell groups more selectively increase their firing rate during SWRs, consistent with previous observations ^38,54,125,126^. In our population, inhibitory cell groups exhibit diverse temporal dynamics around the ripple peak. We also found a small number of ripple-suppressed neurons, primarily among the interneuron groups. In rodents, OLM and axo-axonic cells under anesthesia ^11^, and a subset of axo-axonic cells in behaving rats ^127,128^ decrease their firing rate during ripples (though the OLM class activation depended on the preparation ^28^.) These ripple-suppressed inhibitory cell types can play a special role in long-term synaptic plasticity ^129^ and cell assembly modifications ^31,130,131^, making them of special interest for further study in primates.

Other spike-LFP dynamics included spike-field coherence (SFC) that was strongest at low (<10Hz), beta2/low gamma (25-35Hz), and high gamma (∼80Hz) frequency ranges, consistent with previous findings from macaque hippocampus ^36,132,133^. Cell groups had stronger phase-locking to high-gamma and ripple frequencies compared to beta/low gamma similar to the phase locking reported in mice ^28^. Spike-phase timing in the weaker, mid-band frequencies of 7-30 Hz, spanned the greatest range of phases (∼90°) and exhibited the most groups whose firing phases differed from the pyramidal cells’ phase (Figure S2C). Greater phase variance may signify a more adaptable oscillatory regime that the circuit may capitalize upon during learning, compared to a narrower phase “bandwidth” indicating lower-information-bearing entrainment, as seen in the ‘peak’ frequency-modulated bands ^134^. The spike-phase coherence was additionally modulated by state, with theta-band coherence more pronounced during sleep, and beta/gamma coherence stronger during the task, which generally aligned with previous studies of SFC in macaques ^36,38,132,133^.

Our findings reveal that the average firing phase relationship across cell groups changes dynamically in response to network oscillations. This alteration in relative spike timing may result in the induction of different forms of long-term plasticity ^135^. Additionally, hippocampal sub-circuits may be dynamically organized by phases of local network oscillations to process different spatiotemporal aspects of experiences ^124^. Therefore, our findings of oscillation-dependent relative spike timing may be important for segregating different types of circuit operations for sculpting distinct behavioral-cognitive states. Future research should explore whether and how this variability in phase dynamics within the primate hippocampus correlates with diverse behavioral states.

### Cell type contributions to disease states

Diverse neuron types across the brain cooperate in time to generate network oscillations, regulate synchrony, select cell assemblies, and implement brain states ^24^; thus, it is not surprising that impairments in the activity or loss of specific cell types significantly contribute to a range of disease states and cognitive deficits, from neurodegenerative diseases like Alzheimer’s to psychiatric disorders such as schizophrenia ^25,33–35^. Differences in function among cell types imply that their selective impairment can lead to distinct dysfunctional states. Indeed, subpopulations of pyramidal cells differ in their sensitivity to insults ^12^, and their involvement in generating abnormal activity^136^. Similarly, different populations of hippocampal inhibitory GABAergic interneurons display differential vulnerability to neurodegeneration in Alzheimer’s disease and are preferentially affected at different disease stages ^137^. Another example is mouse models of temporal lobe epilepsy in which CCK-expressing basket cells innervating the perisomatic compartment of the CA1 pyramidal cells are selectively reduced while parvalbumin-containing boutons are conserved ^138^. Thus, understanding the diverse cell types in the brain and their functional organization is crucial for elucidating the mechanisms underlying neurological and psychiatric disorders and developing effective treatments. Currently, this understanding relies almost exclusively on detailed observations in rodent models. Species divergence in GABAergic cell type ^139,140^, however, may lead to the emergence or prevalence of disease states that are specific to primates^141^. One example relevant to memory disorders is that rodents do not naturally develop measurable pathological features of Alzheimer’s disease such as Aβ plaques or p-tau tangles ^142^. As a result, the rhesus macaque monkey is a vital translational animal model for studying disease mechanisms^143^. The current study provides an important step towards the identification of those cell types in the primate hippocampus that may be targeted in these disease models.

### Sublayer-specific regulation of circuit function

One of the most compelling cases for distinct subcircuits within CA1 comes from the relatively recent discovery of superficial/deep pyramidal cell populations, along with their specific inhibitory cell type cohort ^144^. Similar to findings in rodents ^17,22^, macaque CA1 pyramidal cells in superficial and deep layers differ in average firing rate, bursting, and coefficient of variation. These subpopulations of pyramidal cells differed in their pairwise cofiring with several of the inhibitory cell groups. This included a temporally structured opposing interaction between superficial and deep pyramidal cells with ripple-suppressed inhibitory cells, where deep-pyramidal cells were suppressed, in the context of increased superficial-cell responses, presumably arising from somatodendritic inhibitory mechanisms known to differently target pyramidal cells by depth ^13,15,104,105^. Inhibitory-cell switching can determine the precisely-timed expression of independent cell types or assemblies, creating parallel channels for information transmission ^131,144–146^. Evidence for parallel channels can be seen in the biased participation in cell assemblies, which are composed of either superficial or deep pyramidal cells but rarely both ^22^. When added to differences in connectivity, this enables both layer-biased processing of different elements of behavioral tasks and selective coordination of extrahippocampal activity ^22^. We discovered that primate CA1 also demonstrates this cell assembly bias by strata. Because extra-hippocampal projections are organized by the depth of CA1 pyramidal neurons in macaques^147,148^, separable hippocampo-cortical networks may be organized through parallel channels within primate CA1. These findings, together with strata-specific differences in experience-dependent plasticity and in memory formation ^15,21,22,149,150^, suggest that their detection in primates may prove to be key to understanding how the hippocampus structures activity during memory formation.

## Acknowledgments

The funding for this study was provided by National Institute of Neurological Disorders and Stroke Grant (R01) 1RF1NS127128-01, NEI (P30 EY008126), NIH OD (1S10D021771) to the VUIIS Center for Human Imaging, and the Whitehall Foundation Grant 100001391. Declaration of interests: The authors declare no competing interests.

## Methods

### Subjects and Behavioral Design

Two adult female macaques (*Macaca mulatta*, referred to as ‘M1’ and ‘M2’) were subjects in this study. Both monkeys underwent training in a 3D testing enclosure, which allowed them to move freely. This enclosure was equipped with multiple touchscreens distributed around its periphery. To receive a fluid reward, the monkeys needed to move sequentially to each of four touchscreens placed in one corner of the environment. On each screen, they had to touch designated objects associated with that screen, avoiding distractor objects. Their overall performance in completing the four-screen sequence determined their reward. The screens were arranged in a 2×2 array on opposite corners of the 3D space, necessitating visual search, reaching, and walking or climbing during each trial. The monkeys completed various trial blocks, which took place in both screen array corners of the testing apparatus. Following their training session, the monkeys were returned to their housing. For monkey M2, overnight recordings started within approximately 20 minutes of the training session. For monkey M1, there were occasionally 1-2 hour gaps. Sleep epochs occurred with the monkeys in their normal housing area, with their usual social housing accommodations, in complete darkness, following the automated lighting system’s 12/12 overnight dark cycle. Throughout the manuscript, we used the terms “task” and “sleep” to broadly refer to recordings in the 3D testing enclosure and overnight occupancy in the housing, respectively. As a consequence, occasional brief periods of immobility during training were included in the task epoch and initial quiescent waking in the dark, prior to sleep, may have been included in the sleep epoch. All procedures were conducted in accordance with the approved protocols and authorized procedures under the local animal care authorities (Institutional Animal Care and Use Committee).

### Electrophysiological recordings

Active multichannel probes were inserted into a chronically implanted base^37^, including a 128-channel probe (DA128-1, linear configuration with 40μm contact spacing) for Monkey M1 and a 64-channel probe (organized into 4 parallel shanks with 40 channels at 90 μm spacing and 3 shanks with 8 channels at 60 μm spacing) for Monkey M2 (’Deep Array’ design, by Diagnostic Biochips, Inc). The probes were affixed to adjustable microdrives (M1, M2: NanoDrives, Cambridge Neurotech, Inc.; M1: custom, Rogue Research, Inc.) to facilitate precise depth positioning adjustments post-implantation, and allowing the raising and relowering (max. 7 mm or 5 mm, respectively) into target areas while remaining implanted. Post-operatively, the probes were incrementally advanced through these drives in 125mm steps until the target positions were achieved. The localization of recording sites was verified through postoperative CT scans, coregistered with pre-operative MRI data, and also by referencing functional landmarks that changed with increasing depth. Notably, the emergence of depth-specific sharp-wave ripples (SWRs) within unit-dense layers served as a key reference point. To align the 4 shanks in M2, we employed cross-correlation analysis of LFP signals observed during ripple activity across the channels. Local field potentials (LFPs) were digitally sampled at a rate of 30 kHz using the FreeLynx Wireless Acquisition system (Neuralynx, Inc) and subsequently bandpass filtered within the 0.1 Hz to 7500 Hz range. During task performance and sleep recordings for Monkey M2, data were wirelessly transmitted to the Freelynx acquisition system (Neuralynx, Inc). To optimize battery life, sleep recordings for Monkey M1 were stored on an SD card within the acquisition system. A high-frequency noise signal at approximately 6 kHz marked the point at which the battery capacity reached 10%. The onset of this noise was detected by calculating the root mean square (RMS) envelope of the band-passed filtered signal between 5800-6200 Hz. The noise initiation point was defined as the timestamp at which the RMS envelope exceeded 110% of the median value, and sleep data following this point was excluded. For the purpose of merging task and sleep recordings, data was initially converted to microvolt units, bitVolts were standardized to 0.195, and the data was subsequently transformed into binary files encoded as 16-bit integers. Sessions that exhibited no signal loss during recording were exclusively included in the analyses. In LFP-related analyses, the raw signal was subjected to third-order Butterworth filtering with a low-pass cutoff frequency set at either 350 Hz or 450 Hz, and the data was downsampled to 1 kHz.

### CT-MRI image processing and coregistration

The General Registration tool, Elastix, in the Slicer (version 4.11), was used to perform the registration of post-operative CT scans with pre-operative MRI images. The registration process maintained default parameters. Preceding registration, the CT images were cropped near to and including the cranium.

### Power spectral parametrization and fitting

To estimate the layer-specific spectral content of the hippocampal and neocortical recordings, we used Welch’s method with a 50%-overlapping 1024-sample sliding Hanning window to estimate power spectra for the frequency range of 1–150 Hz with a frequency resolution of 0.25 Hz. To identify spectral peaks and compare between task and sleep states, we parameterized power spectra using the method described by ^151^. This method models power spectra as a combination of the 1/f frequency components (aperiodic) in addition to a series of Gaussians that capture the presence of peaks (periodic components). The model was fit to a frequency range between 1 Hz and 200 Hz with a frequency resolution of 0.5 Hz. Settings for the algorithm were set as: peak width limits: (0.5, 12); max number of peaks: infinite; minimum peak height: 0; peak threshold: 2.0; and aperiodic mode: ‘Fixed’.

### Detecting hippocampal sharp-wave ripples and estimating ripple slope

A single hippocampal LFP channel with largest ripple amplitude was selected for ripple detection. The wide-band signal was band-passed filtered between 100-180 Hz using a 3^rd^ order Butterworth filter, and squared signal was calculated. The squared signal was further band-passed filtered in 1-20 Hz range and z-normalized. SWR peaks were detected by thresholding the normalized squared signal at 3SDs above the mean, and the surrounding SWR start and stop times were identified as crossings of 1 SDs around this peak. SWR duration limits were set to be between 40 and 400 ms. Adjacent events with an inter-ripple interval of <20 ms were merged.

#### Exclusion Criteria

In addition to assessing amplitude and duration, we used several criteria to identify and exclude potential false-positive ripple events.

#### Channel Noise Exclusion

A ‘noise’ channel, defined as one devoid of detectable sharp-wave ripples (SWRs) in the local field potential (LFP), was designated. Any events simultaneously detected on this channel were considered as potential false-positives, likely originating from artifacts such as electromyography.

#### Spectral Analysis

We conducted spectral analysis of each detected ripple event to further scrutinize its characteristics. The data was spectrally decomposed using Morlet wavelets, allowing us to compute the frequency spectrum for each event. This was achieved by averaging the normalized instantaneous amplitude within ±50 ms of the ripple peak over the frequency range of 50-200 Hz, normalized by multiplying the amplitude by the frequency. We then analyzed the number and properties of spectral peaks in each detected ripple frequency spectrum. These peaks were identified using the findpeaks function in MATLAB, considering parameters such as peak height, prominence, peak frequency, and peak width. This analysis aimed to ensure that the detected ripple events genuinely reflected high-frequency, narrowband bursts within the ripple band range. We applied multiple criteria to achieve this: first, authentic ripple events were expected to exhibit a predominant spectral peak within the ripple band range. Therefore, if no single prominent peak (corresponding to the ripple band) was identified, the event was rejected. Additionally, authentic ripple events were anticipated to display a limited narrowband burst; thus, if the ripple-peak had an excessively wide peak width (indicating more broadband spectral changes) or prominent high-frequency activity, the event was considered for rejection (see^152^).

#### Visual Inspection

Finally, all detected ripples underwent visual inspection, and any events flagged as false-positives during this process were subsequently removed from the dataset.

#### Ripple slope and power

To estimate ripple slope, we first computed an average ripple signal per recording channel by aligning ripples to their peak. These averages were then smoothed using a moving average method (40ms window). For baseline normalization, we subtracted the value at −50ms from the peak from each channel’s average. Ripple slopes were determined by integrating values from −50ms to 0ms relative to the peak, and these slopes were further smoothed across channels using a 2-channel moving average window. Additionally, for ripple amplitude analysis, we calculated the sum of the absolute values of the average ripple between −50ms and 50ms for each channel, providing a measure of the overall strength of the ripple signal within this time window.

### Spike sorting

Spike sorting was performed using Kilosort 1.0 (^153^, https://github.com/cortex-lab/KiloSort). The process involved applying a 300-Hz high-pass filter to the raw signals, followed by whitening the data in blocks of 32 channels. Parameters relevant to automated sorting are detailed in the table.

#### Removing putative double-counted spikes^154^

The Kilosort algorithm will occasionally fit a template to the residual left behind after another template has been subtracted from the original data, resulting in double-counted spikes. Such double-counted spikes could artificially inflate inter-spike interval (ISI) violations for a single unit or create erroneous zero-time-lag synchrony between neighboring units. Consequently, spikes with peak times within a 5e-4 second interval and peak waveforms detected on the same channels were systematically removed from the dataset.

#### Removing units with artefactual waveforms

Kilosort1 generates templates of a fixed length (2 ms) that matches the time course of an extracellularly detected spike waveform. However, there are no constraints on template shape, which means that the algorithm often fits templates to voltage fluctuations with characteristics that could not physically result from the current flow associated with an action potential. The units associated with these templates are considered ‘noise’ and are removed on the basis of spread (waveform appears on many channels), and shape (e.g. no peak and trough or sinusoidal waveform) criteria and autocorrelogram function.

#### Manual curation and re-clustering with Phy

Manual curation and re-clustering were performed using Phy (https://github.com/kwikteam/phy). Kilosort-derived clusters were imported into Phy for manual curation. Units that were poorly isolated according to the initial Kilosort results were re-clustered using Klusta with custom-designed plugins (https://github.com/petersenpeter/phyplugins) to obtain well-isolated single units. The quality of these clusters was evaluated based on refractory period violations and Fisher’s linear discriminant metrics. Noise clusters and poorly isolated units were subsequently excluded from the analysis.

### Kilosort Parameters

|  |  |
| --- | --- |
| ops.datatype | 'dat' |
| ops.fs | 30000 |
| ops.Nfilt | 1024 |
| ops.whitening | 'full' |
| ops.nSkipCov | 1 |
| ops.whiteningRange | 32 |
| ops.criterionNoiseChannels | 0.2 |
| ops.Nrank | 3 |
| ops.nfullpasses | 6 |
| ops.maxFR | 20000 |
| ops.fshigh | 300 |
| ops.ntbuff | 64 |
| ops.scaleproc | 200 |
| ops.NT | 32*1024+ ops.ntbuff |
| ops.Th | [4 10 10] |
| ops.lam | [10 30 30] |
| ops.nannealpasses | 4 |
| ops.momentum | 1./[20 400] |
| ops.shuffle_clusters | 1 |
| ops.mergeT | 0.1 |
| ops.splitT | 0.1 |
| ops.initialize | 'fromData' |
| ops.spkTh | -4 |
| ops.loc_range | [3 1] |
| ops.long_range | [30 6] |
| ops.maskMaxChannels | 5 |
| ops.crit | 0.65 |
| ops.nFiltMax | 10000 |

### Localization of neuronal somata within the CA1 layers

To estimate the location of the linear-array channels relative to CA1 layers, we used features of the sharp wave ripple. The sharp wave component of the SWR arises from CA3 Schaffer collateral inputs that generate a current sink in stratum radiatum (SR) with a return source centered in stratum pyramidale^155^. In the relatively closed fields of rat or mouse CA1, this is evident as a polarity reversal between SR and the deeper layers, where the envelope flattens, distorts and ultimately reverses^6^. The source/sink gradient has been used as a depth reference to identify superficial/deep pyramidal cells^22^, and should be more sensitive to local generators (more tolerant of open-fields) than the raw fields^156–158^. On this basis, we first estimated the radiatum/pyramidale transition using current source density (CSD), the second spatial derivative, of ripple-trigger averaged signal across the regularly-spaced LFP channels. This identified the general regions of SR and SP. Next, we calculated ripple power and the slope of sharp-wave envelope peak, both of which have been used to center the pyramidal layer in previous studies^17,22^, respectively. For each session, we set the channel closest to the LFP slope zero-crossing as 0, and the depth of the other channels in relation to that point, considering the fixed inter-electrode distance. No additional scaling was made to the inter-channel distances. This reversal point fell in the middle of sinks and sources of CSD and above the depth of maximum ripple power. For the side-shank linear sites (in M2), we adjusted the physiological depth based on the correlational similarity of mean sharp-wave ripple LFP. For all isolated units, the site with the largest spike amplitude for each unit was regarded as the location of the cell body.

### Classification of cell groups

To establish the feature space for cell group classification (i.e. as putative cell types), we leveraged single-unit waveforms and interspike interval (ISI) distributions of the cells. The filtered single-unit waveforms (comprising 48 samples at 30 kHz) were first normalized within the range of 0 to 1 and aligned. Subsequently, we used the Uniform Manifold Approximation and Projection (UMAP) to reduce the dimensionality of the waveform matrix to 2 components. In a parallel process, the log10 ISI distributions within the range of 0-10 seconds were subjected to UMAP analysis to obtain 2 components as well. For these UMAP procedures, we employed a custom UMAP function implemented in MATLAB^159^. The selection of UMAP parameters was guided by consultation with^58^. To generate the feature space for clustering, the four attributes (two from waveform and two from ISI) were concatenated.

### UMAP Parameters

UMAP Parameters:
|  |  |
| --- | --- |
| **min_dist** | 0.1 |
| **n_neighbors** | 20 |
| **n_components** | 2 |
| **n_epochs** | 5000 |
| **metric** | euclidean |

We used Spectral Clustering in MATLAB ^160,161^ to perform the clustering, with Mahalanobis distance serving as the primary distance metric, *k*-means for clustering, and *K* number of clusters where *K* is estimated from the GMM model described below. Spectral Clustering begins by constructing a similarity matrix that encapsulates the pairwise similarities between data points. From this matrix, a graph is formed where data points are nodes connected by edges weighted by their similarities. Thus, clustering is treated as a graph partitioning problem. The Laplacian matrix of this graph is then computed, and its eigenvalues and eigenvectors are used to transform the data into a lower-dimensional space. This process leverages the eigenvectors corresponding to the smallest non-zero eigenvalues to achieve a partition that minimizes within-cluster dissimilarities and maximizes between-cluster differences. Spectral clustering excels at identifying complex cluster structures that might not be linearly separable, making it a powerful tool for grouping data in multidimensional spaces. Compared to GMM, spectral clustering does not assume that the data is grouped in a Gaussian or elliptical manner making it superior for identifying complex structures. As described, spectral clustering works well with data where the clusters are connected but not necessarily compact or isotropic as GMM assumes. It uses graph-based techniques which can capture the “connectivity” of the data points. In addition, spectral clustering allows for the easy incorporation of different similarity/distance measures.

To estimate the number of clusters in the data in an unsupervised way, we used the expectation-maximization (EM) algorithm for Gaussian mixture model (GMM) clustering. We modeled the data as a weighted sum of multivariate Gaussians:

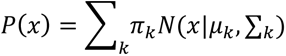

with *k* components parametrized by mean *μ*_*k*_, covariance ∑_*k*_ and mixing coefficient *π*. The EM algorithm fits this model by iteration of a two-step process: it first estimates posterior probabilities of the data given the current set of parameters (E step), and then updates the parameters to maximize the log-likelihood function of the model given the current estimates (M step). The steps are repeated until convergence. We used MATLAB function *fitgmdist* to fit the GMM with initial parameters: *RegularizationValue* = 0.01; *SharedCovariance*: tested with both true and false; *MaxIter* = 10000; *CovarianceType*: tested with both ‘diagonal’ and ‘full’. To select the number of Gaussian components in the model we used the Bayesian information criterion (BIC):

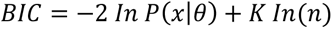

where *P*(*x*|*θ*) is the maximized likelihood for the estimated model, *K* is the number of parameters, and *n* is the sample size. By including a penalty term that grows with the number of parameters, the BIC cost function effectively favors simpler models and reduces overfitting. The optimal number of clusters was chosen as the value that minimized the BIC computed between 1 and 20 components. The GMM+BIC method systematically yielded a larger number of optimal clusters (*k* = 14) compared to the Eigengap heuristics method for different choices of distance metric. The Eigengap method ^161^, while widely used for estimating the number of clusters in spectral clustering, has several limitations including the assumption of cluster size balance, dependence on graph construction, and sensitivity to noise. Thus, we relied on the GMM+BIC to inform the initial number of clusters in the spectral clustering with post-clustering refinements. After initial clustering, we noticed a high level of similarity among some of the clusters. To decide whether to merge or keep split, we combined the waveform and ISI attributes of the members of the similar clusters and created a GMM model from those with the same parameters as described but the K was chosen to be between 1:2. Based on the results of the BIC, we decided to keep them as separate if K=2 was the optimal number of clusters, otherwise, we merged the clusters. The 10 resultant clusters are after these stepwise refinements.

### Classification of deep and superficial CA1 pyramidal cells

The first group was designated the pyramidal cell group, because these cells demonstrated the low firing rates, high propensity for bursting, and dense localization within the Stratum Pyramidale that are the hallmarks of CA1 pyramidal cells^53,60,61^. For the superficial and deep analyses, we removed from this group the spatial outliers (with relative depths falling outside the 5-95th depth percentile range), and cells with mean firing rates > 1 Hz or a burst index exceeding 3, before calculating the median depth of these units to segregate into superficial and deep categories. The spatial distribution of CA1sup neurons spanned from −300 to 540 μm, and for CA1deep neurons, it extended from −630 to −330 μm.

### Burst Index

We calculated a burst index to capture the propensity of neurons to discharge in bursts (i.e. short intervals, or “complex spikes”). This was estimated from the mean of spike auto-correlogram (1 ms bin size) measured between 1 and 10 ms, normalized by the mean value between 200 and 300 ms^104,162^.

### Ripple-associated spike content analysis

Within-SWR firing rate was calculated as the number of spikes during SWRs divided by the cumulative duration of the SWRs between the first and last spike fired by the cell. The ripple participation probability of individual units was defined as the fraction of SWRs in which that neuron fired at least one spike. Ripple ratio was defined as firing rate during ripple events divided by the overall firing rate.

To calculate spike density functions, spike vectors of individual cells were binned with 1 ms binsize and peri-event time histograms (PETHs) were computed locked to ripple peaks and converted to rate. Then a gaussian kernel with a 10 ms S.D. was applied to the PETHs to obtain spike densities. For each cell, we calculated the baseline activity as the mean firing rate of a shufled surrogate dataset created for that cell and subtracted this value from the original PETH.

To test the hypothesis that spiking activity of single cells are modulated surrounding the ripple peak, we derived nonripple event surrogates. These surrogates were created for individual cells using timestamps selected randomly without replacement from recording epochs when ripples were not detected. We matched the number of ripple and nonripple surrogate events (n observed ripple events = n nonripple events). Furthermore, to ensure that signal properties were maximally matched between target events and surrogates, surrogates were drawn only from a 10-min time window before and after the corresponding ripple event. To test the significant difference between the ripple-locked and surrogate spike densities, we used a cluster-based permutation procedure using 5000 permutations and a cluster threshold of p < 0.05 and a final threshold for significance of p < 0.05.

### Spike-field synchronization

To quantify spike-field synchronization, we used the pairwise phase consistency (PPC) measure, which is unbiased by the number of spikes^163^. For hippocampal recordings, we selected the sharp-wave ripple channel in the Stratum Pyramidal of the CA1 across sessions. If the unit spikes were from this channel, we used an adjacent channel to measure the PPC. The spectral content was estimated with Morlet wavelet decomposition method using a constant number of cycles (7) per frequency for frequencies between 1 and 200 Hz with a frequency resolution of 1 Hz. We measured PPC values separately for spikes during task and sleep states, and only for neurons that fired at least 100 spikes^163^. One caveat of this approach is that it assumes signals to be oscillatory and, additionally, sinusoidal. This can distort preferred phase estimations. We used an alternative approach to estimate phase of modulation in restricted time windows of presumed oscillatory bouts, across frequency ranges. For each frequency group, LFP signal was filtered in the specified frequency band using a 3^rd^ Butterworth filter. Next, instantaneous phase and power were derived from Hilbert transform. A phase value was assigned to each action potential during significant periods (power of filtered LFP > 2 S.D. above the mean) using linear interpolation. Peaks are at 0° and 360° and troughs at 180° throughout the paper. Then we calculated the mean phase and resultant vector using the circular statistics toolbox on MATLAB^96^ (https://github.com/circstat/circstat-matlab). To find the grand mean phase and resultant vector of a cell group in a specific frequency, we measured the mean phase of firing for included members of that cell group and then calculated the mean phase and resultant vector of the mean phases. The grand mean phase and resultant vector determines the consistency of phase of firing among the members of a cell group. For the calculation of the grand mean phase, we included only cells that had at least 20 spikes during significant epochs of interest. We set a significance threshold of p = 0.01 using the Rayleigh test for phase locking.

### Pairwise cofiring

We used two complementary approaches to estimate pairwise cell interactions. First, recordings were segmented into 25 ms time bins, and for each neuron, the number of spikes within each bin was counted and converted to firing rate. Subsequently, we convolved the spike vectors with a Gaussian kernel, with a standard deviation calculated as binsize / (2 * sqrt(2 * log(2))) (converted to Full Width at Half Maximum, FWHM). Pearson’s correlation coefficients were calculated between the spike density vectors of different neurons, serving as a measure of their co-firing tendencies.

To prevent global periods of inactivity or instability from influencing the Pearson’s coefficients, we only considered correlations if two conditions were met: 1) the overlapping windows of activity exceeded 5 minutes, and 2) both cells fired at least 100 spikes within the overlapping time window.

Additionally, cross-correlograms (CCGs) were computed between pairs of neurons with a 1 ms bin resolution and converted to rate (CCG divided by the reference cell’s number of spikes per time bin). These CCGs were smoothed using a zero-lag partially hollowed Gaussian filter with a convolving window of 5 ms and a hollow fraction of 0.6 ^105,164^.

The mean activity in a baseline window (−50 to −20 ms) was subtracted from the original CCGs to selectively evaluate the fine temporal interactions (+/-20 ms). To assess the statistical significance of CCGs, we created surrogate spike time series’ by shufling the inter-spike intervals of the cells ^125^ and calculating their CCGs. This process was repeated 5000 times to generate a surrogate distribution of CCGs. P-values were computed for each time bin of the CCG (−20 to 20 ms) and subjected to multiple-comparison correction using the Benjamini and Hochberg False Discovery Rate (FDR) procedure To compare CCGs between the CA1sup and CA1deep groups, a two-sample permutation test (n=5000) with Tmax correction was used ^165^.

### Assembly pattern identification and activation strength

Cell assemblies were identified during sleep recordings as previously described^22,166–170^. Significant co-firing patterns were detected using an unsupervised statistical method based on independent component analysis (ICA). The spike trains for each neuron were binned into time windows of 90 ms (corresponding to the 90^th^ percentile of ripple durations) and z-score transformed to eliminate biases due to differences in average firing rates. This creates for each session, a cell x binned firing rate matrix (Z). Next, a principal component analysis was applied to the matrix (Z). For this, the correlation matrix of Z was given by 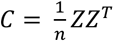 and the eigenvalue decomposition of C was given by:

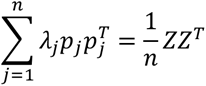

where *λ*_*j*_ is the j^th^ eigenvalue of C and *p*_*j*_ is its corresponding eigenvector. The Marcenko-Pastur law was used to estimate the number of significant patterns embedded within Z. For a nXB matrix, an eigenvalue exceeding *λ*_*max*_, defined by 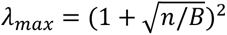, signifies that the pattern given by the corresponding principal component explains more correlation than would be expected if the neurons were independent of each other. The number of eigenvalues exceeding *λ*_*max*_ was defined as NA and therefore represents the minimum number of distinct significant patterns in the data. The significant principal components were then projected back onto the binned spike data

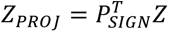

where *P_SIGN_* is the nXNA matrix with the NA principal components as columns.

Independent component analysis (ICA), using the fast ICA algorithm (http://research.ics.aalto.fi/ica/fastica), was then applied to the matrix ZPROJ. That is, an NAXNA unmixing matrix W was found such that the rows of the matrix *Y = W^T^Z_PROJ_* were as independent as possible. The arbitrary signs of the independent component (IC) weights were set so that the highest absolute weight was positive. The unmixing matrix W was then used to derive each cell’s weight within each assembly *V = *P*_SIGN_W* where the columns of *V* (*i. e., v_1_, …, v_NA_*) are the weight vectors of the assembly patterns.

To determine the strength of the expressed assemblies, we tracked each assembly pattern *v*_*k*_ over time by:

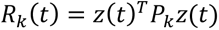

where *z*(*t*) is a smooth vector function containing for each neuron its z-scored instantaneous firing rate and *P*_*k*_ is the matrix projecting *z*(*t*) to the activation strength of the assembly pattern *k* at time *t*.

### Organization of cell assemblies

The majority of the identified assembly patterns had a characteristic distribution whereby a few neurons displayed high weights, while a larger group of neurons had weights approximating zero. To determine the significant membership corresponding to each assembly pattern, we used the criterion that the member neurons of an assembly should have weights exceeding the mean weight by at least two standard deviations. All subsequent analyses were conducted directly on the assembly patterns themselves, using the weight vectors derived from the contribution of all recorded neurons. Based on the composition of significant assembly members, the assemblies were classified into three distinct groups. **Within Assemblies:** These assemblies had at least one significant member from the pyramidal cell group. If all significant pyramidal members were exclusively from the superficial or deep regions, we labeled the assembly as “within superficial” or “within deep,” respectively. **Across Assemblies:** These assemblies included at least one significant member from each of the superficial and deep pyramidal cell groups. **Non-pyramidal cell assemblies:** This category included assemblies that didn’t have any significant pyramidal cell members.

To estimate the probability of realization of a specific assembly organization, we used the binomial probability density function (binopdf in MATLAB). Observed and expected probabilities were computed as:

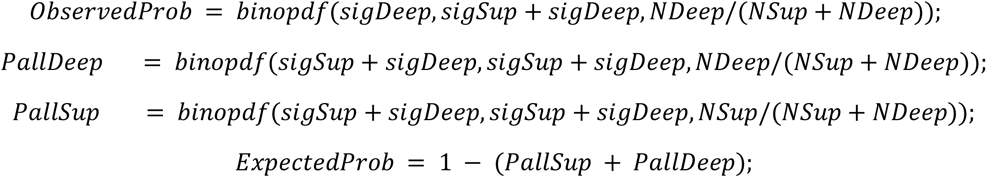

Where *sigDeep* and *sigSup* represent the number of significant CA1deep and CA1sup members within the assembly, respectively. *NDeep* and *NSup* denote the total number of recorded CA1deep and CA1sup neurons within the session. *ObservedProb* signifies the probability of encountering the specific organization of the assembly. *PallDeep* and *PallSup* indicate the probability that all significant pyramidal members exclusively belong to either CA1deep or CA1sup. *ExpectedProb* reflects the anticipated probability that the assembly could belong to the “across” category.

Assembly participation probability was calculated for all cells per session and it was defined as the number of assemblies where a cell participated as a significant member divided by the total number of detected assemblies in a session.

### Statistical analysis

Data collection was conducted prior to (thus blind to) the clustered cell groups. The sharp wave ripple and general spiking activity were the main features determining any change made to electrode depth. Data analysis and behavioral experiments did not require manual scoring. No specific methodology was used to estimate the minimum required population sample, due to no knowledge of the (cell group, assembly) population sizes during data collection; however, the number of animals and sessions and the general registration of unit activity across channels was comparable to or greater than those used in prior studies. All statistical analyses were performed using MATLAB R2021a, utilizing non-parametric methods for comparisons of means and variances, including Kruskal-Wallis analysis of variance, two-sample permutation tests, one-sample randomization tests.

#### Two-sample permutation test

A two-sample permutation test is a statistical method used to compare two independent groups or samples in a hypothesis testing framework. It is particularly valuable when the data do not meet the assumptions of parametric tests like the t-test. For comparison in which each sample had a single data point, we shufled the membership assignments, computed the mean difference between the surrogate samples, and repeated this process 5000 times to create a surrogate probability distribution of mean differences. The original, non-permuted data are then compared to the surrogate distribution to obtain uncorrected p-values.

For *cluster-based multiple comparison correction*, all samples with p-values smaller than 0.05 were selected. These selected samples were subsequently clustered into connected sets based on their adjacency, and the size of each cluster was calculated. This process was repeated 5000 times to generate a distribution of cluster sizes. Clusters with sizes exceeding the cluster threshold at the 95th quantile were reported as significant ^171^.

For CCG analyses we controlled family-wise error rate (FWER) using the Tmax correction method ^165^. This method provides strong control of FWER, even for small sample sizes, and is much more powerful than traditional correction methods ^172,173^. It is also relatively insensitive to differences in population variance when samples of equal size are used ^174^.

One-Sample Randomization Test: This test, akin to the permutation test, involved comparing a time series against a surrogate dataset created from randomly selected timestamps.

For all other posthoc tests, either Tukey-Kramer or the Benjamini & Hochberg multiple comparison correction was applied, as specified in the main text.

Boxplots were used to present data, with the median, 25th, and 75th percentiles represented within the box, and the whiskers illustrating the data range. In cases where boxplots did not display individual data points, outliers were excluded from the plots but were consistently included in the statistical analysis.

---

**Figure S1:**
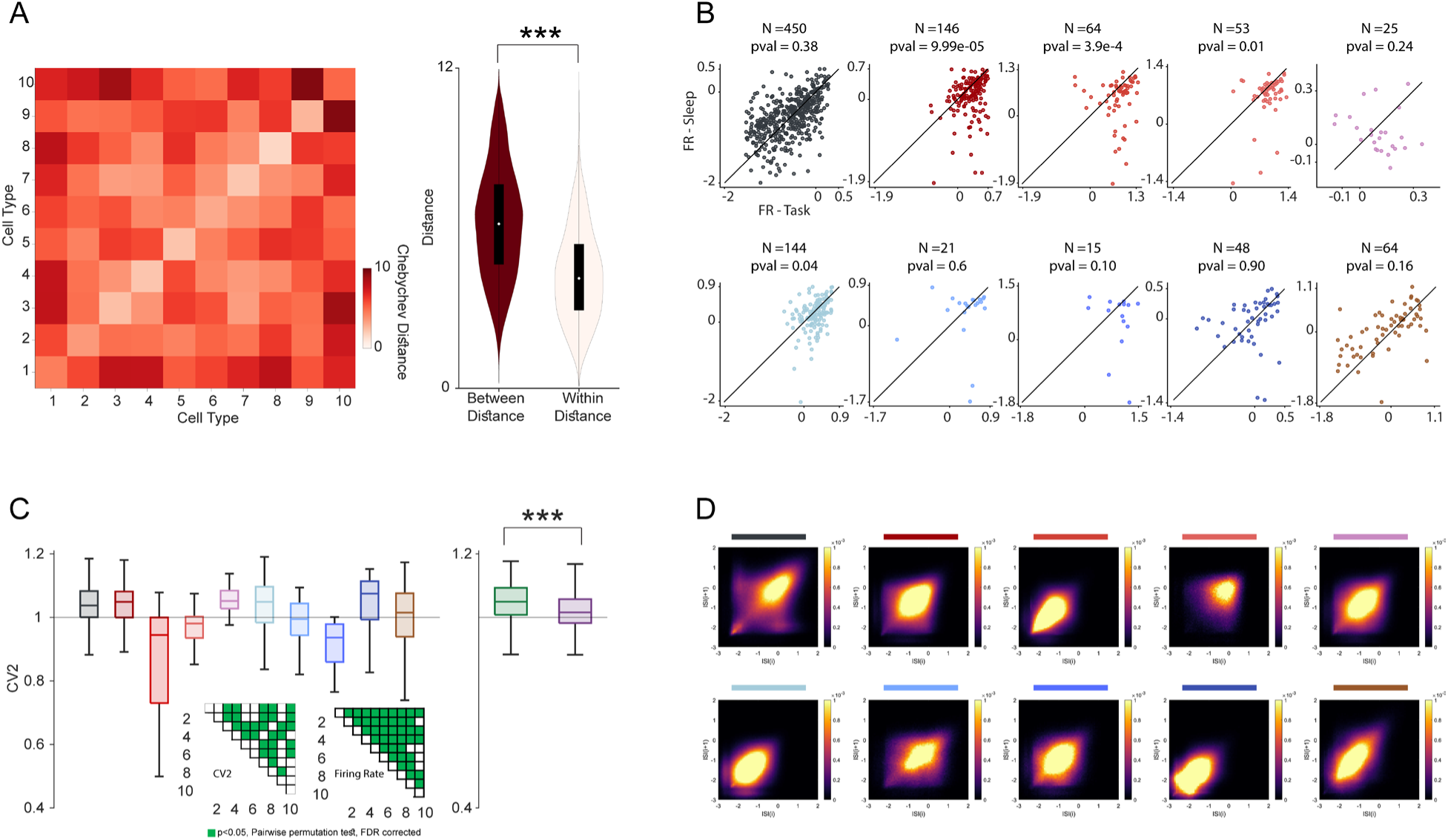
Physiological features of cell groups in CA1. **(A)** Left: Chebyshev distance matrix in feature space, for cells according to assigned cell group. Note that the diagonal values are smaller than off-diagonal, indicating that within-group members were closer together in the feature space compared to between-group members. Right: Distribution of between-group (dark red), and within-group (bone white) of distance values (***p < 0.001, two-sample permutation test; dots indicate medians and black bars quartile ranges) **(B)** Comparison of firing rates during task compared to sleep states, organized by cell group. The color of the data points indicated the cell group membership. (N = number of cells; pval = result of a two-sample permutation test). **(C)** Left: Box plot distributions of ISI-based coefficients of variance (CV2) for each cell group. Right: CV2 for superficial versus deep pyramidal cell groups (p<0.001, two-sample permutation test). Inset: the left matrix shows the result of pairwise permutation tests following significant Kruskal-Wallis test, p < 0.001; dark pixels indicate p < 0.05). The right matrix shows the same for firing rate, for comparison. **(D)** Joint ISI histograms. Each figure shows the distributions of spike intervals at the next interval (ISI_i+1_) as a function of the current interval (ISI_i_), averaged across cells of a given cell group. Horizontal bar colors indicate cell group membership.

**Figure S2:**
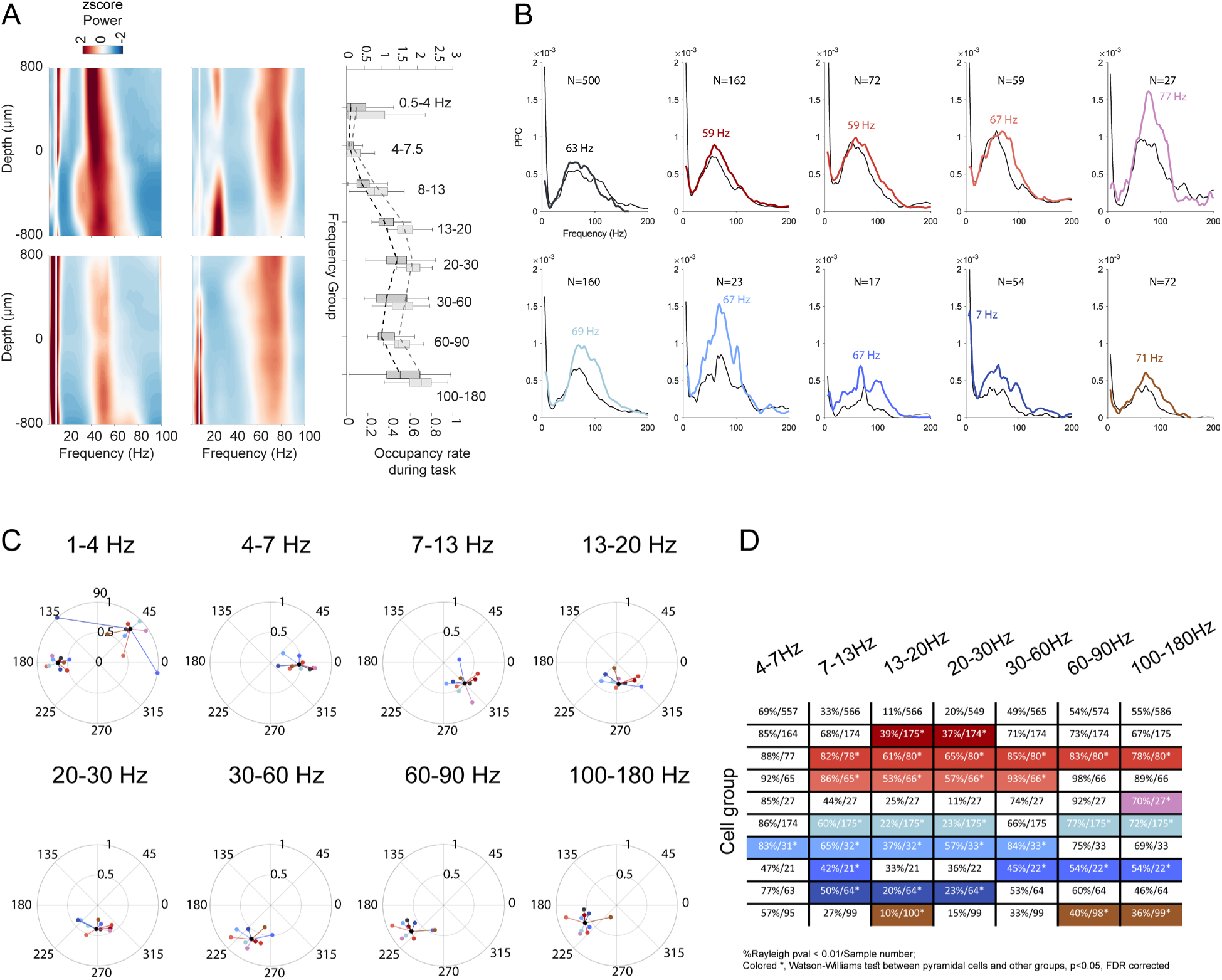
Spectral profiles and spike-field relationships. **(A)** Left: 1/f corrected FOOOF power spectrum across depths of recording in CA1 during task (top) and sleep (bottom) for each animal (M1, left; z-normalized per channel). Right: Corrected (black, top axis) and balanced (grey, bottom axis) proportion of detected oscillatory bouts during task compared to sleep states across frequency bands (i.e., prevalence). LFP channels were selected from Stratum Radiatum. **(B)** Mean PPC values during task (colored) and sleep (grey). The peak frequencies during task are listed. **(C)** Spike phase-of-firing plots separated for each frequency band. Each plot shows, for the specified frequency band, the grand mean phase (black) and distance to each cell group’s resultant vector, taken from distribution of the underlying cells’ resultant vectors. Note that for the 1-4 Hz group, we separated the results for the 2 animal subjects due to conspicuous differences between them, unique to this plot. **(D)** The matrix shows the results of pairwise permutation tests between the pyramidal cell group and other groups, for each frequency band (p < 0.05, two-sample Watson-Williams test, FDR corrected)

**Figure S3:**
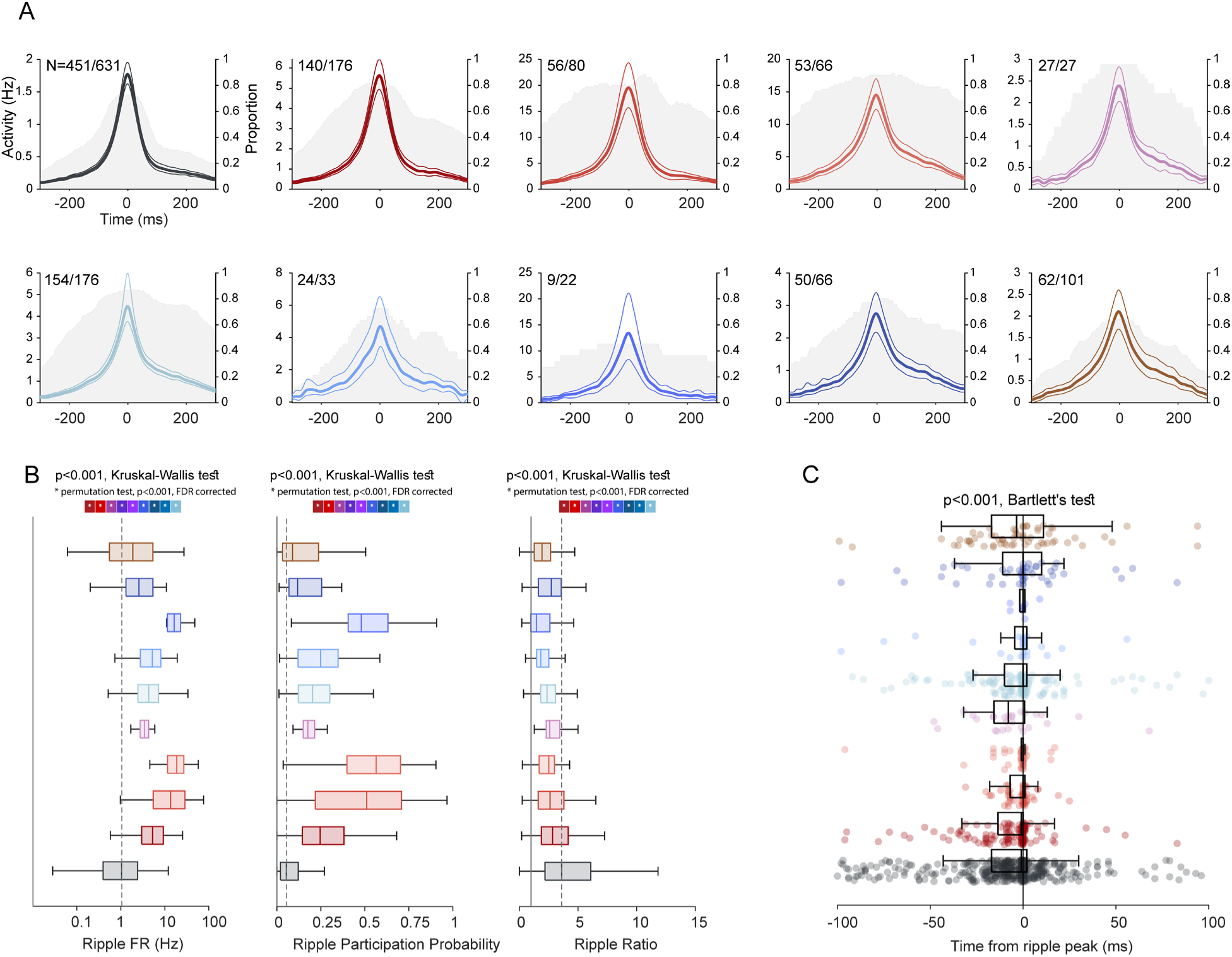
Sharp-wave ripple-associated spiking dynamics. **(A)** Ripple-aligned spike density (mean + 95% CI) for significantly modulated cells of different groups. N: Significantly modulated/total cells in the group. Proportion of significantly modulated units are shown in the shaded area plots, with proportions listed on the right axis (p < 0.05, one-sample cluster-based permutation test). **(B)** Left: Ripple firing rate across cell groups; dotted line indicates PYR group median. Middle: Ripple participation probability. Right: Ripple ratio (average firing rate during ripples / overall firing rate) by cell group, with ratio = 1 indicated by black line. Dotted line indicates PYR group. All plots show (p < 0.001, Kruskal-Wallis test) and asterisks in color coded box show the result of post-hoc pairwise permutation tests between the pyramidal cell group and all other cell groups (* p < 0.001, FDR corrected). **(C)** The distribution of peak response times around the ripple peak, separated by cell group. (p < 0.001, Bartlett’s test for equality of variances).

**Figure S4:**
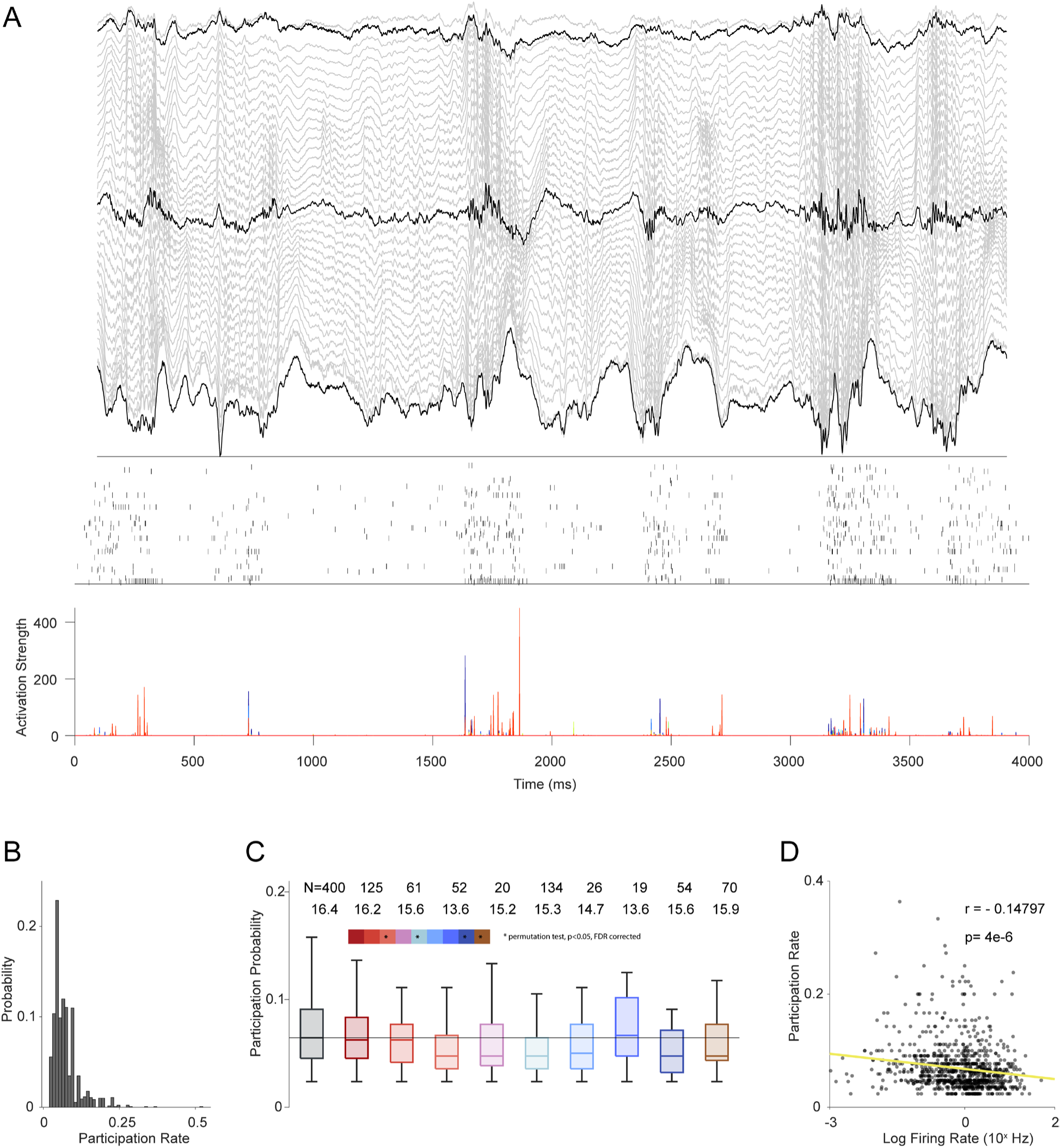
Organization of assembly dynamics. **(A)** Top: Example LFP traces across depth of recording in CA1 during sleep. Middle: Raster plot of simultaneously recorded units. Bottom: Assembly activation strength. Each assembly is assigned a different color. **(B)** Distribution of assembly participation rate for each cell group. Cells with a participation rate of 0 were removed from the analysis. **(C)** Distribution of assembly participation probability for each cell group (p < 0.001, Kruskal-Wallis test). The first row of numbers shows number of recorded units that participated in at least one assembly. The second row of numbers reports the average number of assemblies available to participate in, for each cell group, across session. The thin black line indicates the pyramidal cell median. The color-vector shows the post-hoc pairwise permutation test between pyramidal cell group with all other cell groups. (* p < 0.05, FDR corrected) **(D)** Relationship between log firing rate of neurons and assembly participation rate, revealing an inverse correlation (r = −0.14, p < 0.001 t-test).

